# Comparison of Wild Type DNA Sequence of Spike Protein from SARS-CoV-2 with Optimized Sequence on The Induction of Protective Responses Against SARS-Cov-2 Challenge in Mouse Model

**DOI:** 10.1101/2021.08.13.456164

**Authors:** Sheng Jiang, Shuting Wu, Gan Zhao, Yue He, Jiawang Hou, Yuan Ding, Linlin Bao, Jiangning Liu, Chuan Qin, Alex Cheng, Brian Jiang, John Wu, Jian Yan, Ami Patel, David B. Weiner, Laurent Humeau, Kate Broderick, Bin Wang

## Abstract

COVID-19 caused by SARS-CoV-2 has been spreading worldwide. To date, several vaccine candidates moved into EUA or CA applications. Although DNA vaccine is on phase III clinical trial, it is a promised technology platform with many advantages. Here, we showed that the pGX9501 DNA vaccine encoded the spike full-length protein-induced strong humoral and cellular immune responses in mice with higher neutralizing antibodies, blocking the hACE2-RBD binding against live virus infection in vitro. Importantly, higher levels of IFN-γ expression in CD8+ and CD4+ T cell and specific cytotoxic lymphocyte (CTL) killings effect were also observed in the pGX9501-immunized group. It provided subsequent protection against virus challenges in the hACE2 transgenic mouse model. Overall, pGX9501 was a promising DNA vaccine candidate against COVID-19, inducing strong humoral immunity and cellular immunity that contributed to the vaccine’s protective effects.

## INTRODUCTION

Severe acute respiratory syndrome coronavirus 2 was a member of the *Coronaviridae* family, which caused Coronavirus Disease 2019(COVID-19)^1,2^. Over 103 million confirmed cases of COVID-19, including 2 million deaths, were reported to WHO (as of February 4, 2021, https://covid19.who.int/)). SARS-CoV-2 is a single-strand positive-sense RNA virus constituted of Spike protein, Nucleocapsid, Membrane protein, and Envelope protein^3^. The spike protein was primarily associated with the infection ability of the virus and constituted of two distinct subunits, namely S1 that was composed of the receptor-binding domain (RBD), and S2 consisting of the transmembrane domain. The two subunits were activated after host cell cleavage and bound to human angiotensin-converting enzyme 2 (hACE2) for cell entry ^4^. Therefore, the spike protein was an ideal immunogen candidate. The neutralization antibody which blocks RBD binding to hACE2 inhibited virus infection^5^, and the mutation of amino acid of spike protein affected the infectivity and stability of virus^6^.

Technologies to make a vaccine against the COVID-19 includes inactivated virus, subunit protein, mRNA, or viral vector. DNA vaccine technology plays an essential role in vaccines. Compared to other vaccine technologies, DNA vaccine is easier and cheaper to make, thermally stable induces both cell-mediated and humoral immune response ^7–9^. Prior works demonstrated that DNA vaccines could induce higher neutralizing antibody responses against viruses such as SARS, Middle East respiratory syndrome (MERS), Zika virus^10–13^.

DNA vaccine can also induce strong T cell immunity than other vaccines. During virus infection, T cell immunity plays an important role that CD4^+^ T cells are thought to provide necessary secondary signals like cytokines and inflammatory signals to promote immunity, and at the same time, CD8+ T cell express effector genes including Granzymes and perforin to enhance its cytotoxic capacity to direct kill virus^14^. However, the magnitude of T cell immunity in being beneficial or harmful for SARS-CoV-2 patients was unclear. Therefore, we test our DNA vaccine T cell immunity and explore its role in virus infection in the hACE2-transgenic mouse model.

Here, we demonstrate that the pGX9501 DNA vaccine candidate, optimized codon sequence for the full-length spike protein from the SARS-CoV-2, showed a significant level of neutralizing antibodies and robust cellular responses and resulting in complete protection in a challenge animal model.

## METHODS AND MATERIALS

### Animal experiment

Female, C57BL/6 mice (6-8 weeks of age) and Balb/c mice (6-8 weeks of age) were purchased from Beijing Vital Laboratory Animal Technology Co., Ltd. (Beijing, China) and Shanghai Jiesjie Laboratory Animal Co., Ltd. (Shanghai, China), which were kept in SPF condition. hACE2 transgene BALB/c mice were from the Institute of Laboratory Animal Sciences, CAMS&PUMC. All animal experiments were approved by the Experimental Animals Committee of SHMC.

The mice were injected twice via the intramuscular route (i.m.) with 25μg plasmid, and electroporation was followed at intervals of two weeks. Serum was collected 14 days after the second immunization.

### Plasmid preparation

Plasmids of pGX9501, pVAX1-S-WT, and pVAX1 were transformed into DH5a E. Coli, respectively. A single colony was undergone expansion in a one-liter flask for culturing in LB broth. Plasmids were extracted and purified by MaxPure Plasmid EF Giga Kit (Magen, China). The final purified plasmids were dissolved in saline buffer at 1mg/ml. The purity was measured by an agarose gel electrophoresis and a UV detector at a range of 1.8-2.0 OD260nm/280nm. Endotoxin in those plasmids was below 30 EU/mg by LAL test.

### Rare codon analysis

The sequence of wild type and the sequence optimized pGX9501 was submitted to the GenScript Rare Codon Analysis Tool (https://www.genscript.com/tools/rare-codon-analysis). From origin organism to expression host, the tool compared the difference between sequence by analyzing the DNA sequence features like cis-regulatory and negative repeat elements that could influence transcription and translation efficiencies after construct entered is unclear.

### Antigen-specific humoral immune responses

Enzyme-linked immunosorbent assay (ELISA) was used to measure the antigen-specific antibody production induced by the DNA vaccinations as previously described. Briefly, 96-well plates were respectively coated with 0.5μg/ml of pre-S1 (Sino biological, 40591-V05H1), 0.5μg/ml of pre-S2 (Sino biological, 40590-V08B), and 0.17ug/ml of RBD (Sino biological, 40592-V08B) protein (50 mM carbonate-bicarbonate buffer, pH 9.6) at 4°C overnight and blocked with 5% BSA in PBST (0.05% Tween 20 in PBS) at 37°C for 1 hour. The plates were incubated with diluted serum from different immunization groups for 1 hour at 37°C. Antibodies were detected with HRP-conjugated goat anti-mouse IgG (Southern Biotech, Birmingham, AL) after the enzymatic reaction was developed, the OD values were read at 450/620 nm by an ELISA plate reader (Bio-Rad, Hercules, CA).

### hACE2-RBD blocking assay

The blockade of hACE2 binding to SARS-CoV-2 RBD in ELISA as previously reported^15^. Briefly, 96-well plates were coated with 0.34μg/ml of RBD protein (50 mM carbonate-bicarbonate buffer, pH 9.6) at 4°C overnight and blocked with 5% BSA in PBST (0.05% Tween 20 in PBS) at 37°C for 1 hour. Plates were incubated with diluted serum samples from different immunization groups for 1 hour at 37°C. Then, 0.12μg/ml of hACE2 (Sino Biological, 10108-H08H) was added and reacted for 1 hour at 37°C. The Goat anti-Rabbit IgG-HRP (Invitrogen, 32460) and Rabbit anti-hACE2 antibody (Sino Biological, 10108-RP01) were used to detect the concentration of hACE2 binding with the RBD. The following formula calculated the blocking ratio: Blocking Ratio= (1-(experimental group/control group)) *100%.

### Live virus neutralization assays

Neutralization assays were performed at the Institute of Laboratory Animal Sciences, CAMS&PUMC of China. Seed SARS-CoV-2 (SARS-CoV-2/WH-09/human/2020) stocks and virus isolation studies were performed in Vero E6 cells, which are maintained in Dulbecco’s modified Eagle’s medium (DMEM, Invitrogen, USA) supplemented with 10% fetal bovine serum (FBS), 100 IU/mL penicillin, and 100 μg /mL streptomycin, and incubated at 36.5 °C, 5% CO2. Virus titers were determined using a standard 50% tissue culture infection dose (TCID50) assay. Serum samples collected from immunized animals were inactivated at 56 °C for 30 min and serially diluted with a cell culture medium in two-fold steps. The diluted samples were mixed with a virus suspension of 100 TCID50 in 96-well plates at a ratio of 1:1, followed by 2 h incubation at 36.5 °C in a 5% CO_2_ incubator. 1× 10^4^ Vero cells were then added to the serum-virus mixture, and the plates were incubated for 3–5 days at 36.5 °C in a 5% CO_2_ incubator. Cytopathic effect (CPE) of each well was recorded under microscopes, and the neutralizing titer was calculated by the dilution number of 50% protective condition.

### Flow cytometry

Single suspension cells from spleens and lymph nodes collected 14 days after the second immunization were prepared and stimulated with 10μg/ml peptides at 37°C for 5h. Cells were stained with viability dye eflour780 in PBS for 15 minutes on ice followed by washes twice with PBS supplemented with 2% FBS. For the detection of cell surface antigens, cells were stained with fluorochrome-tagged antibodies, as shown in the following table for 15 minutes on ice. To detect intracellular cytokines or intranuclear transcription factors, cells were fixed and permeabilized using an intercellular cytokine staining kit (BD) or a commercial transcription factor staining kit (eBioscience). All stained samples were run on LSRFortessa (BD) and analyzed by FlowJo (TreeStar).

| Antibody | Company | Clone | Lot number |
| --- | --- | --- | --- |
| anti-Mouse CD4-APC | ebioscience | GK1.5 | 4329627 |
| anti-Mouse CD8a-PerCP/Cy5.5 | biolegend | 53-6.7 | B219152 |
| anti-Mouse TNFα-PE | ebioscience | MP6-XT22 | 438513 |
| anti-Mouse Granzyme B-PE Cyanine7 | ebioscience | NGZB | 4281151 |
| anti-Mouse IFNγ-APC | ebioscience | XMG1.2 | 4289683 |
| anti-Mouse IFNγ-BV421 | biolegend | XMG1.2 | B232596 |
| anti-Mouse CD3e-FITC | ebioscience | 145-2C11 | E00061-1632 |
| anti-Mouse IL-5-PE | ebioscience | TRFK5 | 12-7052-82 |
| anti-Mouse IL-13-eFlour710 | ebioscience | eBio13A | 46-7133-82 |

### SARS-CoV-2 Spike protein Peptide pool

COVID-19 spike RBD peptide pool (SARS-CoV-2 Spike protein 258-518aa) published previously^16^ was used for the study.

### Cytotoxic lymphocyte (CTL) killing ability

Singles suspensions of splenocytes from naïve syngeneic mice were diluted to 1.5*10^8^/ml by RPMI1640 with 10% FBS and 2% Penicillin and Streptomycin, respectively then pulsed with or without 5ug/ml peptides as mentioned above at 37°C. After 4 hours, a higher concentration of eflour450 (ebioscience, 65-0842-85) at 5mM was used to label pulsed peptide cells. Cells without peptide-pulsed were labeled with a low concentration of eflour450 at 0.5mM at room temperature in the dark. After being rinsed by PBS three times, 4*10^6^ of labeled and peptides-pulsed cells and another equal number of labeled cells without peptide-pulsed were adoptive transferred by tail vein injections into mice previously immunized with different vaccines, respectively. Six hours later, the percentage of labeled cells was detected with LSRFortessa flow cytometry (BD) and analyzed by FlowJo (TreeStar). The following formula calculated the specific cell lysis: Specific cell lysis ability= (1-(percentage of cells incubated with peptide/percentage of cells incubated without peptide)) *100%

### SARS-CoV-2 challenge Study

SARS-CoV-2 (SARS-CoV-2/WH-09/human/2020/CHN) was isolated by the Institute of Laboratory Animal Sciences, CAMS&PUMC. Immunized hACE2 transgenic mice were infected with SARS-CoV-2 (10^6^TCID_50_) via the nasal route in a volume of 100μl 7 days after the final immunization. Five days after infection, the Lung was harvested for measuring virus loads by qPCR and H&E staining. Before the challenge, serum was collected for ELISA to evaluate the neutralizing antibody levels.

### Statistical Analysis

The statistical analysis methods and sample sizes (n) are specified in the results section or figure legends for all quantitative data. All values are reported as means ± sem with the indicated sample size. No samples were excluded from the analysis. All relevant statistical tests are two-sided. P values less than 0.05 were considered statistically significant. All animal studies were performed after randomization. Statistics were performed using GraphPad Prism 7 software. In all data *p<0.05, **p<0.01, ***p<0.001, and ****p<0.0001.

## RESULTS

### Expression Level of Optimized Spike Sequence

A DNA construct, pVAX1-S-wt, made from the wild type sequence of the full-length spike protein of the SARS-CoV-2 was subcloned into the pVAX1. The sequence of the same region was optimized via SynCon^®^ technology as previously described, synthesized, and cloned into pVAX1 as the pGX9501^16^. Two constructs were transfected into 293T cells in vitro parallelly under the same condition and subjected into a qRT-PCR and Western blotting Analysis to compare expression levels. As depicted in Figure 1A and 1B, the level of mRNA and protein expressions were hundred times greater from the optimized pGX9501 over the wild type pVAX1-S-wt. We further discovered that the critical differences between the two sequences were the contents of negative Cis-elements, five in the wild-type, one in the optimized pGX9501 (Table 1).

**Figure 1.**
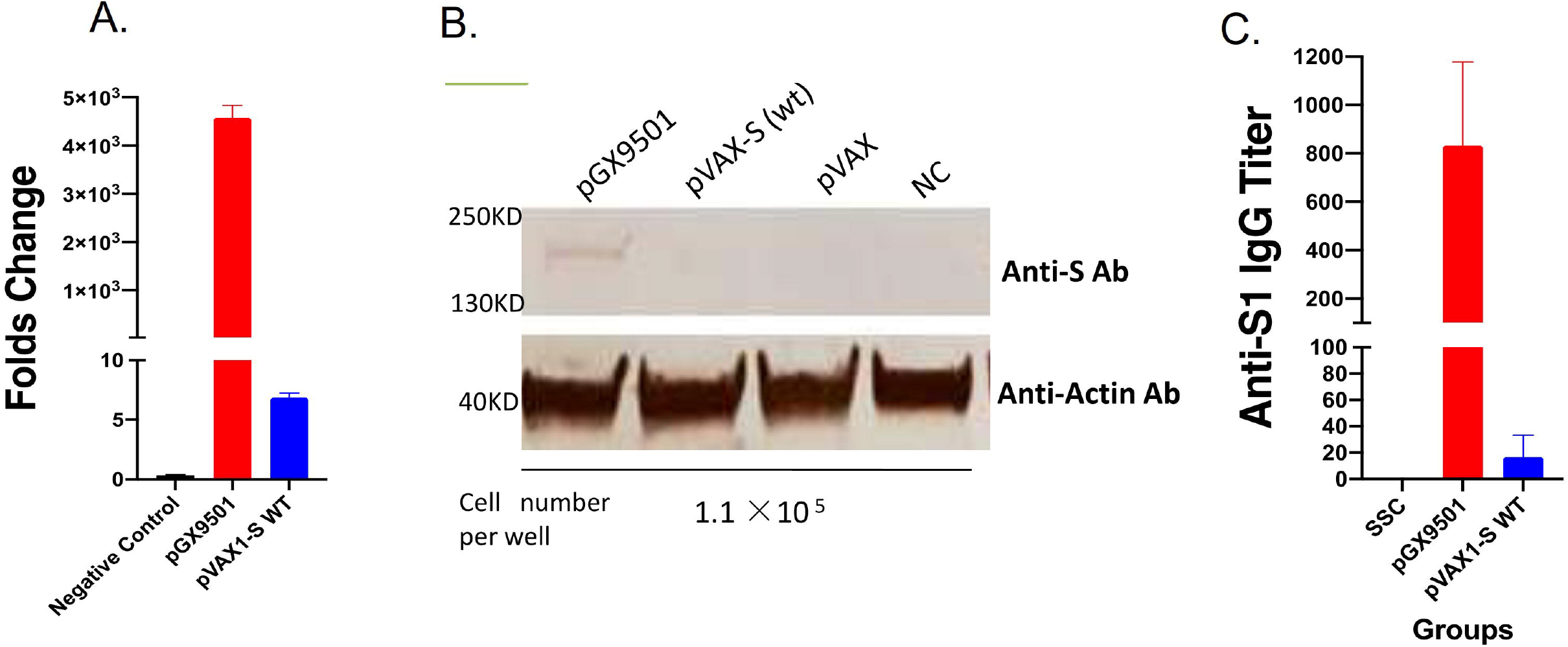
Comparison of expression and antibody levels of optimized versus non-optimized spike sequences. 293T cells were transfected by pGX9501, pVAX1-S-WT, and pVAX1 for 48hrs and lysed for RT-PCR (A) and Western blotting (B), respectively. (C) Balb/c mice were immunized by each construct at 25ug intramuscularly once and antiserums samples 14 days later were used analyzed against S1 recombinant protein in ELISA.

**Table 1.** Sequence optimization score of the optimized and wild type sequences

|  | CAI | GC% | Negative CIS elements | Negative repeat elements |
| --- | --- | --- | --- | --- |
| **pGX9501 Spike DNA** | 0.92 | 53.84 | 1 | 0 |
| **COVID-19 WT Spike DNA** | 0.88 | 51.43 | 5 | 0 |

Notes:
- 1.CAI (codon adaptation index) was to evaluate the optimized sequence based on CIS-regulatory elements, codon usage bias, GC rich and etc. The higher CAI value is better for the optimized sequence. - 2.CIS regulatory elements was including TATA box, termination signal and protein binding sites.

### Effects of Antibody Production and Functional Assay

To evaluate the effects of pVAX1-S-wt and pGX9501 DNA vaccines on the abilities to induce specific antibodies, mice were injected twice at 25μg DNA vaccine each time via intramuscularly (i.m) at biweekly intervals and followed by electroporation by a Cellectra2000 device (Figure 2A). Serum IgG samples were serial diluted and tested for binding titles against recombinant spike proteins covered the receptor-binding domain (RBD), S1 region, and S2 region in the extracellular domain (ECD). Levels of antibodies taken from pGX9501 immunized Balb/c, or the C57/BL6 animals, were a thousand times higher than pVAX1-S-WT immunized (Figure 2B). In addition, these sera were also used to examine the inhibition experiment of RBD to human angiotensin-converting enzyme 2 (ACE2) binding. We observed that a higher level of inhibition was achieved from sera immunized with pGX9501 than pVAX1-S-WT (Figure 2C).

**Figure 2.**
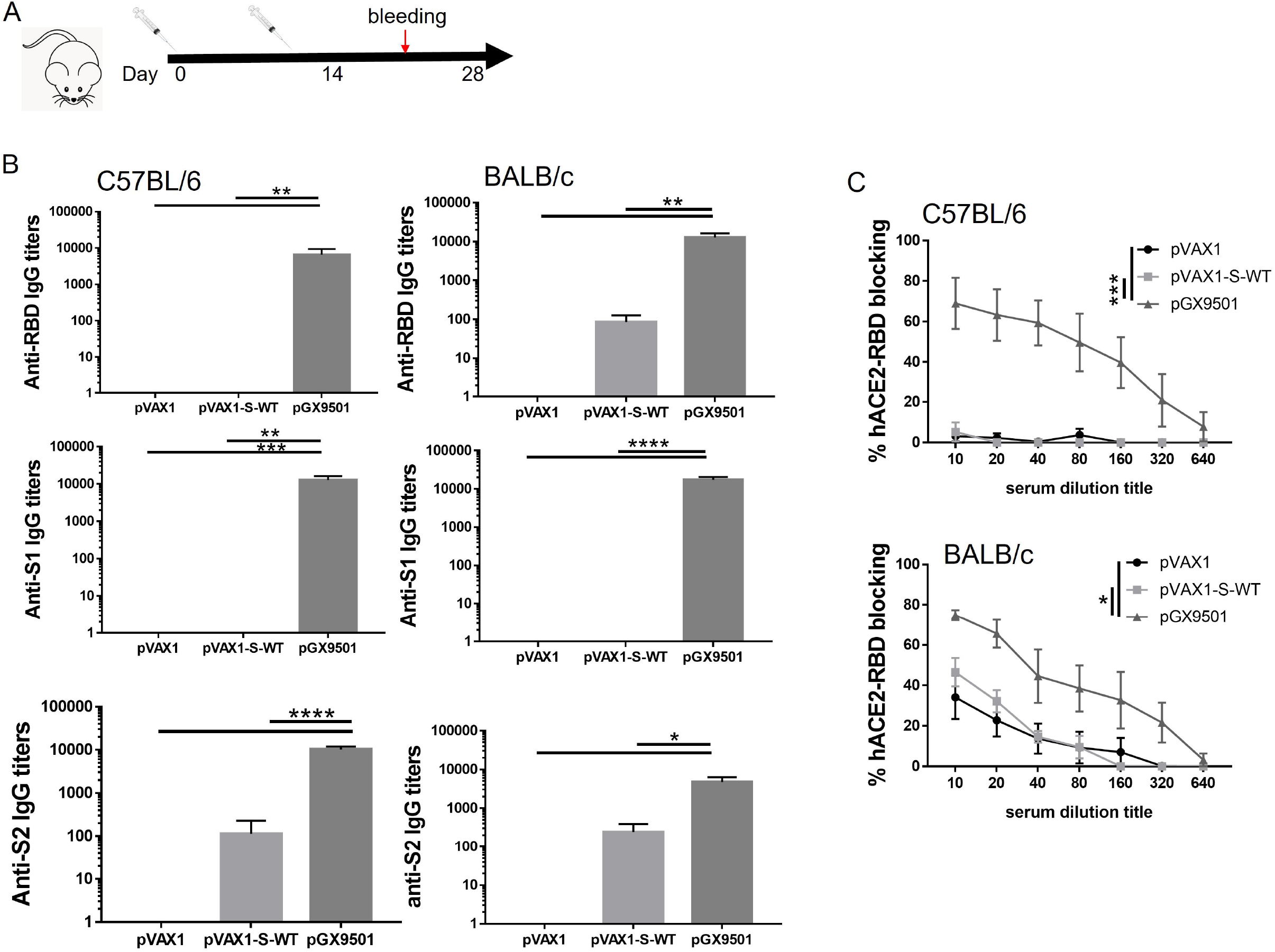
Effects of Antibody Production and Functional Assay. A, The scheme of mice immunizations. B, C57BL/6, or Balb/c mice (n≥ 5 per group) were either immunized with pVAX1 (blue circle) or vaccinated with pVAX1-S-WT (red square) and pGX9501 (green triangle) intramuscularly, following by electroporation. Serum IgG binding titers (mean ± SEM) to SARS-CoV-2 pre-S1, S2, and RBD were measured on day 28. C, Blocking abilities of RBD binding to the hACE2 with serum samples at serial dilutions on day 28. Data shown represent mean blocking efficiency (mean± SEM) for the five mice.

### Efficacy of protective response against SARS-CoV-2 challenge in hACE2 transgenic mice

The goal of a vaccine is to protect against disease, or ultimately against infection of the virus. We examined the efficacy of protection after the DNA vaccine immunization(s) followed by a challenge of SARS-CoV-2 in hACE2 transgenic animals(Figure 3A). After once or twice immunizations of those transgenic mice, animals were challenged intranasally with 1×10^^^6 TCID50 of SARS-CoV-2 viruses post of 7 days from the last immunization. The virus neutralizing assay was performed using serum samples from the pGX9501 immunized once or twice and the samples from pVAX1 as the control. The neutralizing level from pGX9501 immunized once was around 1:18 but increase significantly after the two immunizations at 1:166, compared with the control (Figure 3B). The animals were sacrificed 5 days after the challenge for Analysis of viral loads in lungs and pathological changes. We observed almost 6log_10_ reduction of viral loads in lungs from mice immunized twice with the pGX9501, which was nearly a complete protection against SARS-CoV-2 infection, whereas the 1-2 log10 reductions were reached in animals immunized once with pGX9501 (Figure 3C). According to the pathological Analysis H&E stained lung tissue sections, the mice immunized twice was showed slight infiltration of inflammatory cells in the alveolar septum and pulmonary vessels, compared with the control group (Figure 3D&Table 2).

**Figure 3.**
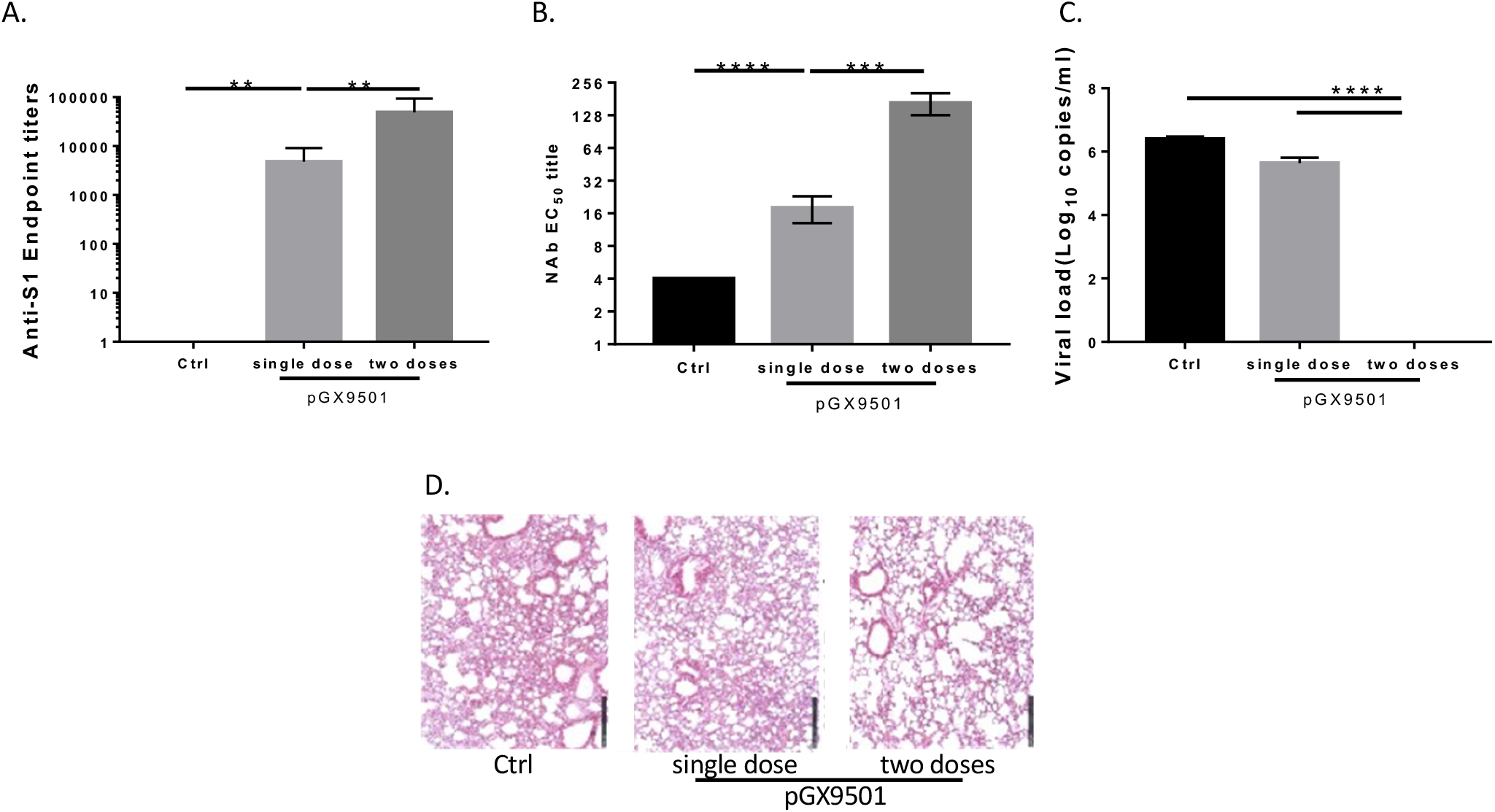
pGX9501 protects against disease challenges with SARS-CoV-2. Mice treated with the vaccine was challenged by SARS-CoV-2 (10^5^TCID_50_) in a volume of 100μl 7days after the second immunization (single dose group was challenged by virus 14 days after immunization). Five days after the challenge, Serum was collected for anti-s1 ELISA(A) and Lung was harvested for measuring virus load by qRT-PCR(B). C, Mice treated with the vaccine was challenged by SARS-CoV-2 (10^5^TCID_50_) in a volume of 100μl 7days after the second immunization (single dose group was challenged by virus 14 days after immunization). Serum was collected for ELISA to evaluate the Neutralizing antibody. D, The histochemistry analysis of lung after H&E staining.

**Table 2.** Analysis of H&E staining of lung of mice challenged with SARS-CoV-2

| Groups | Infiltration of inflammatory cells in alveolar septum |  |  | Infiltration of inflammatory cells around pulmonary vessels |  |  |
| --- | --- | --- | --- | --- | --- | --- |
|  | None | Slight | Medium | None | Slight | Medium |
| 1x SSC (Ctrl) | — | — | ++ | — | + | — |
|  | — | + | — | — | — | — |
|  | — | — | ++ | — | + | — |
|  | — | — | ++ | — | + | — |
|  | — | + | — | — | + | — |
|  | — | — | ++ | — | + | — |
| pGX9501 ( Two dose ) | — | + | — | — | — | — |
|  | — | + | — | — | + | — |
|  | — | + | — | — | — | — |
|  | — | + | — | — | — | — |
|  | — | + | — | — | + | — |
| pGX9501 ( one dose ) | — | — | ++ | — | + | — |
|  | — | + | — | — | + | — |
|  | — | + | — | — | + | — |
|  | — | — | ++ | — | + | — |
|  | — | + | — | — | + | — |
|  | — | — | ++ | — | + | — |
|  | — | — | ++ | — | + | — |

Notes:

### Effects of Cell-Mediated Immunity (CMI)

DNA vaccine has a strong ability to induce CMI due to its expression in host cells and to present its antigens via MHC-I and -II. To assess the CMI, mice immunized with either pGX9501 or pVAX1-S-WT in Balb/c and C57/BL6 mice on day 14 after the second immunization were used to isolate T cells from spleens or Lymph nodes for analyzing intracellular cytokine expressions by flow cytometry. As depicted in Figure 4, percentages of IFN-γ-expressing CD4 T cells, representing Th1 response, were higher in both C57/BL6 and Balb/c mice immunized with pGX9501 than that of the other two groups (Figure 4A&4B). However, expressions of IL-5 and IL-13 in CD4 T cells, representing Th2 response, showed no much differences among the groups (Figure 4C&4D). Levels of IFN-g and TNF-a expressing CD8 T cells again were higher in the group immunized with pGX9501 in both C57/BL6 and Balb/c mice (Figure 5A&5B). Expression of Granzyme B (Gz-B) in CD8 T cells was observed as similar results as obtained as the IFN-g (Figure S1A&S1B).

**Figure 4.**
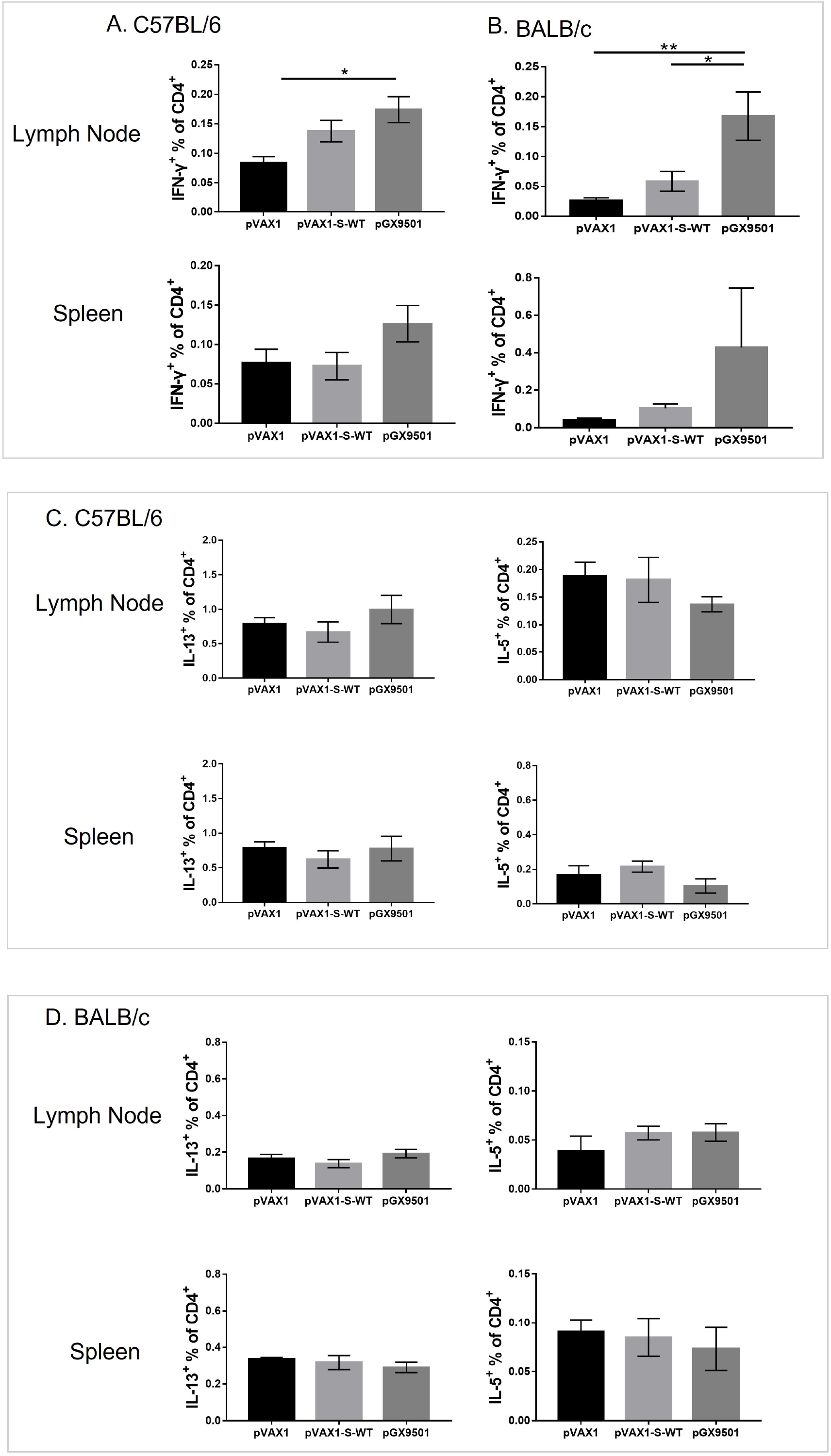
pGX9501 promoted TH1-associated cytokine and did not affect TH2-associated cytokine. Single suspension of splenocytes and lymphoid cells of lymph nodes harvested from C57BL/6 (A) or BALB/c (B) mice immunized were stimulated with 10 mg/mL SARS-CoV-2 peptide pools in vitro for 4 to 6 hours, and IFN-□ production of CD4^+^ T cells was measured by flow cytometry. Both in C57BL/6 and Balb/C mice model. The specific Th2-cytokine expression was verified with the SARS-CoV-2 peptide pool.

**Figure 5.**
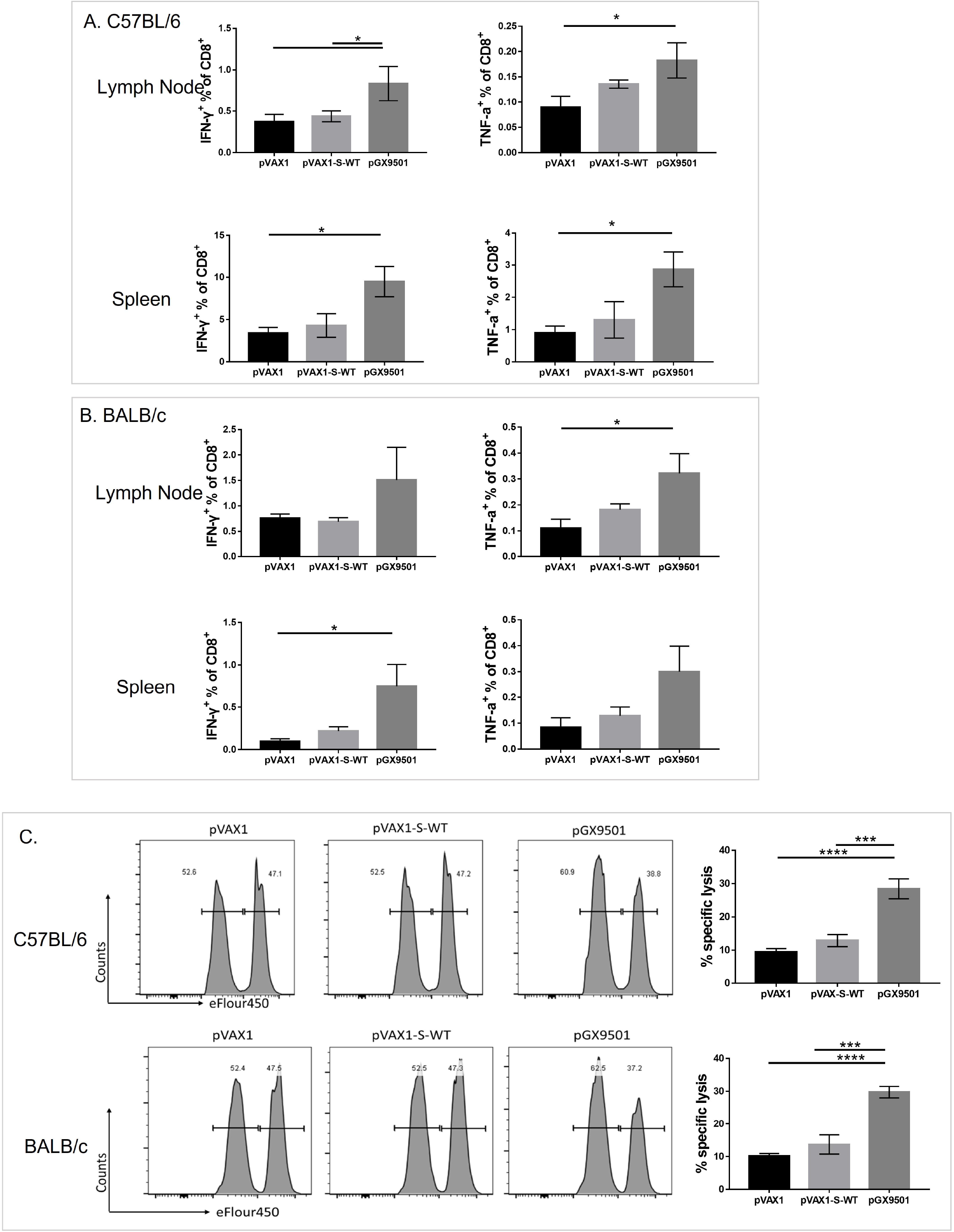
pGX9501 derived more effective specific cytotoxic lymphocyte(CTL) killing ability in vivo and enhanced cytotoxic cytokine expression of specific CD8+ T cell. Single suspension lymphocytes of spleens or lymph nodes from immunized C57BL/6 (A) and Balb/C (B) mice were stimulated with 10 mg/mL SARS-CoV-2 peptide pools in vitro for 4 to 6 hours. Levels of IFN-γ and TNF-α production in CD8+ T cells were measured by flow cytometry. C, Antigen specific cytotoxic lymphocyte(CTL) killing ability was evaluated by an in vivo CTL assay. Target cells at 4*10^6^/ml from naïve mice labelled with a higher concentration of eFlour450 were incubated with 10 mg/mL SARS-CoV-2 peptide pools in vitro for 4-6h before transferring into the immunized mice intravenously. The intensity of eFlour450 peptide labelled target cells was compared with the non-peptide labelled negative control cells after 5 hrs by flow cytometry.

Functional CD8 T cells serve as cytolytic killer T cells to destroy virally infected host cells, resulting in a sterilizing immunity. Since its function can be tested in the in vivo CTL assay, we examined the level of specific CTL killing activities in those immunized groups. We observed that a significantly higher level of CTL was achieved from both Balb/c mice and C57/BL6 mice immunized with pGX9501 compared with that of the other two groups (Figure 5C).

## DISCUSSION

The COVID-19 DNA vaccine (pGX9501) has been found to induce a significantly high level of immunogenicity in animal studies and human trials^16^. Before select candidate constructs for a pre-clinical project, we found that using wild type sequence of SARS-CoV-2 Spike as a DNA vaccine, pVAX-S-wt, induced poor antibody induction. To investigate the reason, we compared those two head-to-heads in this study and observed that over a hundred times higher expression level of mRNA had been achieved with the pGX9501 after transfected into culture cells over that from the pVAX-S-wt (Figure 1A). Such a superior level was also revealed from the corresponding expressed spike protein (Figure 1B). The higher expression level is translated into a higher level of anti-spike antibodies, blocking RBD to hACE2 binding and neutralizing activities against live viruses infecting host cells. Not only was antibody level greatly enhanced, induction of T cells, including CD4 and CD8 T cells producing IFN-g and TNF-a, were also augmented. Notably, the level of cytolytic CD8 cells was significantly increased, demonstrated by the in vivo CTL assay, suggesting the ability to clearance of viral infection. Finally, the codon-optimized pGX9501 construct effectively expresses eukaryotic cells as one of the critical factors, which could be translated into higher efficiency to induce host immune responses, further providing a protective response against live virus challenge with reduced lung pathogenesis in a dose-dependent manner.

Although optimization of codons in a foreign gene could significantly enhance its expression, in general, has been reported, the over hundreds if not thousand hold increases in expression from the pGX9501 to pVAX-S-wt is un-thinkable. Further rare codon analysis observed that five negative Cis-regulatory elements were situated within the coding region of wild-type spike versus only one with the pGX9501^16^ (Figure 1B). Previous studies showed that a gene expression could be inhibited negative cis-regulatory elements and restored after the elements were mutated^17,18^. Thus, we reasoned that the decreasing number of negative CIS-regulatory elements in the optimized pGX9501 could account for the higher mRNA and protein expression over that of wild type besides the rare codon usages.

In SARS-CoV-2 infection, neutralizing antibodies were essential in preventing viral entry and patients’ prognosis. The neutralizing antibody specific to spike protein correlated with a fatal outcome for patients infected with SARS-CoV-2^19^ and prevented viral infection during the viral outbreak for normal individuals^20^. More than 11 anti-spike monoclonal antibodies^21^ including LY-CoV555^22^, 414-1^21^, and CB6^23^ revealed high-affinity bindings to the RBD and hACE2 binding inhibitions correlated to their neutralizing activities. Thus, we also evaluated the specific anti-Spike binding antibody and RBD-hACE2 binding inhibition ability, in which the optimized pGX9501 induced a significantly higher level of the wild type one.

T cell immunity was indispensable for viral clearance demonstrated in animal models infected with viruses like JEV, DENV, and recent Zika^24–29^. In the SARS-CoV infection model, enhanced CD8+ T cell resulted in earlier virus clearance and increase survival^30^. Clinical data in SARS infected patients showed that a better recovery was usually linked with T cell immunity^31^. A similarity has been observed in SARS-CoV-2 infected patients in that COVID-19 patients with non-to mild-symptom showed significantly higher levels of anti-SARS-CoV-2 CD4 and CD8 T cells versus that of the lower level of such T cells in severe or dead patients^32^. Simultaneously, patients infected with SARS-CoV-2 with worse disease outcomes were related to T cell exhaustion^33^, and lymphopenia was accentuated in symptomatic SARS-CoV-2 patients with pneumonia than those without pneumonia that indicated T cell immunity was played an essential protective role in pre-existing immunity against SARS-CoV-2^32,34–36^.

Further supported activations of specific T cells were essential and correlated with protective efficacy by the recently developed mRNA vaccines (BNT162b2 and mRNA-1273)^37^ and adenoviral vector-based one (AZD1222)^38^. In a recent study of AD26.COV2.S Vaccine, CD8 T cell immunity presents a broad spectrum of protection against SRAS-CoV-2 variants^39^. Therefore, antigen-specific T cell responses play an indispensable role against SARS-CoV-2 infectivity and COVID-19 pneumonia. Therefore, T cell immunity is a crucial factor for protecting people from SARS-CoV-2 infection. A DNA vaccine is known to induce higher T cell-mediated response from previous studies, including the pGX9501 induced T cell responses in mice, monkeys, and humans recently in phases I & II. In this study, we observed that mice immunized with the pGX9501 produced a higher expression level of Th1 cytokine IFN-γ and a vigorous CTL activity, which indicate that both Th1 and cytotoxic T cell was involved in the immunity for protecting mice from SARS-CoV-2 infection.

Our study found that the pGX9501 immunized twice showed a better virus clearance and Lung protection than the pGX9501 immunized once. Firstly, the pGX9501 immunized twice induces a higher neutralizing antibody than the group immunized once. Then, the pGX9501 immunized once only showed serval protection effect on virus infection, while the pGX9501 immunized twice protects mice from virus infection. In summary, only one dose may not induce a strong enough immune response to protect the body from the virus, and the pGX9501 immunized twice can. For the substantial virus clearance of the pGX9501 immunized twice group, we thought that neutralizing antibody that inhibits the virus entry and cell immunity that clears the virus in the body was indispensable. The neutralizing antibody the pGX9501 immunized twice induced was highly correlated to the inhibition ability of virus infection^19^. At the same time, the mice immunized twice showed a more IFN-γ and TNF-α expression of T cell, which was associated with virus clearance^26,29,34^. In conclusion, compared with the wild-type coding sequence, the optimized one pGX9501 was a promising DNA candidate vaccine for preventing SARS-CoV-2, and T cell immunity plays a primary role in protecting the body from SARS-CoV-2 infection.

## Acknowledgment

This work was supported by the Chinese National Natural Science Foundation (81991492 and 82041039) and National Key R&D Program of China (2018YFC0840402) to B.Wang.

## Competing interests

Authors declare that they have no competing interests.

## Notes

### Competing Interest Statement

The authors have declared no competing interest.

