## supplementary figure 1 for "Comparison of Wild Type DNA Sequence of Spike Protein from SARS-CoV-2 with Optimized Sequence on The Induction of Protective Responses Against SARS-Cov-2 Challenge in Mouse Model"

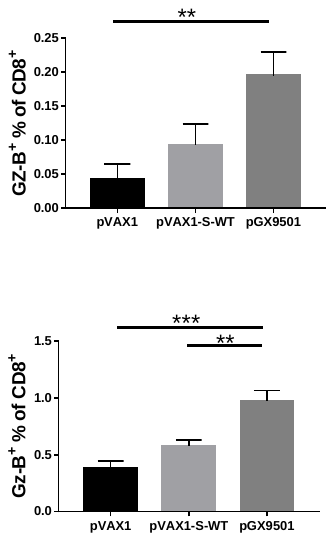

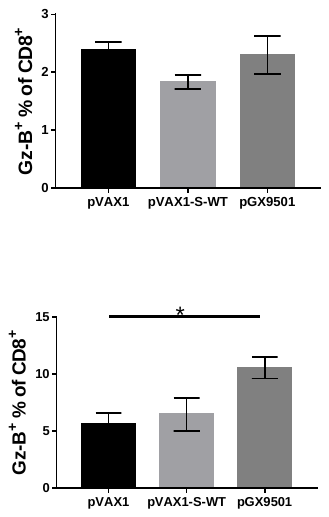

**Supplementary Figure 1.**

A Single suspension lymphocytes of spleens or lymph nodes from immunized C57BL/6 (A) and Balb/C (B) mice were harvested and stimulated with 10 mg/mL SARS-CoV-2 peptide pools in vitro for 4 to 6 hours. The level of Granzyme B production of in CD8+ T cells was measured by flow cytometry.

**Supplementary Figure 1**

A. C57BL/6

Lymph Node

Spleen

B. BALB/c
